# Combined HDAC and ATM inhibition with radiation in preclinical glioblastoma models

**DOI:** 10.64898/2026.10.06.756776

**Authors:** Susan Qualls-Histed, Kaitlyn Chung, Gozde Uzunalli, Sonam Patel, Elizabeth Hayes, Kyria Kelley-Wolfe, Elaine Cabrales, Tanmayee Durshelwar, Liiban Said, Nicole E. Salisbury, Siddhi Desai, Keely Orndorff, Nader Sanai, An-Chi Tien, Artak Tovmasyan, Shwetal Mehta

## Abstract

**Background:** Brain-penetrant histone deacetylase inhibitors (HDACi) are promising radiosensitizing agents for glioblastoma (GBM). HDAC inhibitors can impair DNA damage response (DDR) pathways and promote persistent DNA damage. Here, we evaluated whether combined inhibition of HDAC and the DNA repair kinase, ataxia-telangiectasia mutated (ATM), could synergize with radiation (IR) and improve therapeutic efficacy in GBM.

**Methods:** *In vitro* dose-response assays were performed in patient-derived xenograft (PDX)-derived glioma stem cells (GSCs) with both inhibitors of HDAC (quisinostat, QST) and ATM (AZD1390) with or without IR treatment to assess synergy. Pharmacodynamic responses were evaluated by immunoblotting for histone acetylation, DDR signaling, and DNA damage markers. Transcriptomic analyses were performed to evaluate treatment-associated changes in DNA repair pathways. The combination of QST, AZD1390, and IR was further evaluated in an orthotopic GBM survival study.

**Results:** Combined treatment with QST and AZD1390 produced synergistic radiosensitization in multiple GSC models *in vitro*. Furthermore, QST combined with IR suppressed transcription of homologous recombination (HR) pathway genes in GSCs, impairing double-strand break repair capacity through an epigenetic mechanism that does not directly inhibit ATM kinase activity. Pharmacodynamic analyses confirmed that QST increased histone acetylation while AZD1390 suppressed ATM signaling, together impairing complementary mechanisms to increase radiosensitivity of GSCs. In an orthotopic GBM model, QST combined with IR significantly prolonged survival compared with IR alone. However, the addition of AZD1390 did not provide a significant survival benefit beyond QST+IR, and AZD1390 combined with IR did not significantly improve survival relative to IR alone.

**Conclusions:** Combined HDAC and ATM inhibition with QST and AZD1390 enhances radiosensitivity in patient-derived GBM cells, mechanistically underpinned by QST-mediated increase in DNA damage and dual disruption of DDR through complementary, non-overlapping mechanisms. However, this *in vitro* synergy did not confer additional survival benefit beyond QST+IR in an orthotopic GBM model under the dosing conditions tested, suggesting pharmacological limitations with the tumor microenvironment or suboptimal dosing schedules may have constrained *in vivo* efficacy.

## Background

Glioblastoma (GBM) is the most common and lethal primary brain tumor in adults, with a median survival of 15 months and a 5-year survival rate of only 7% (1). Despite major advances in understanding GBM biology (2), treatment options have remained largely unchanged since the approval of Stupp regimen in 2005 (3). This therapeutic stagnation reflects several persistent challenges, including inadequate drug penetration across the blood-brain barrier (BBB), extensive inter- and intratumoral heterogeneity that drives treatment resistance, and the inability to achieve complete surgical resection (4). Taken together, these challenges underscore the urgent unmet clinical need for novel therapeutic strategies.

Epigenetic dysregulation is one of the key contributors to GBM progression. Histone deacetylases (HDAC) regulate numerous cellular processes, including transcription, differentiation, apoptosis, cell cycle arrest, angiogenesis, metabolism, and the DNA damage response (DDR). As a result, HDAC inhibitors (HDACi) have emerged as a promising therapeutic strategy in cancer (4–7). Several HDACi are already approved for hematologic malignancies and are under investigation in solid tumors, including GBM (8, 9). Among these, quisinostat (QST) is highly potent against HDAC1 and has demonstrated efficacy against patient-derived glioma stem cells (GSCs) both *in vitro* and *in vivo* (10). Although QST significantly extended survival in preclinical GBM models, tumors ultimately recurred, underscoring the need for effective combination strategies.

GSCs contribute to therapeutic resistance in GBM through enhanced DNA repair capacity (11), which enables efficient recovery from therapy-induced damage but this dependency also represents a targetable vulnerability exploitable with DDR inhibitors (12–14). We previously found that QST increases DNA damage in both acutely treated mouse tumors and patient-derived GSCs *in vitro* (*10*), while prolonged treatment suppresses the expression of DDR genes, specifically homologous repair pathway genes. These findings suggest that QST not only induces genotoxic stress but also weakens the repair machinery in GSCs. To exploit this vulnerability, we evaluated QST in combination with AZD1390, a brain-penetrant inhibitor of ATM kinase, a key regulator of the DDR signaling in response to DNA double-strand breaks (DSBs) (14). We hypothesized that combined inhibition of HDAC and ATM would further compromise DNA repair and enhance the effects of IR.

Here, we evaluated the therapeutic efficacy of combining QST, AZD1390, and IR in patient-derived GSC cells and an orthotopic GBM patient-derived xenograft (PDX) model. This triple combination produced a synergistic effect across multiple GSC lines, with significantly elevated DNA damage accumulation. However, ATM inhibition provided no significant survival benefit beyond QST plus IR treatment in an orthotopic GBM model. Collectively, these findings demonstrate that combined HDAC and ATM inhibition can produce robust radiosensitization *in vitro*, however this effect does not translate into improved *in vivo* efficacy. These results provide a preclinical rationale for DDR-targeted combination strategies while also underscoring challenges of translating *in vitro* synergy into meaningful therapeutic benefit *in vivo*.

## Methods

### Cell culture

Patient-derived GSC lines (GB187, GB239, GB126, GB282) were established from resected GBM tumor tissue at the Barrow Neurological Institute. GBM43 cells were obtained from the Sarkaria laboratory at the Mayo Clinic (Rochester, Minnesota, USA). Cells were cultured as previously described (9, 10). Briefly, GSCs were maintained in DMEM/F-12 medium supplemented with B27 and N2 (Invitrogen, Thermo Fisher Scientific), 1% penicillin–streptomycin, Normocin, and growth factors consisting of 20 ng/mL epidermal growth factor (EGF) and 20 ng/mL basic fibroblast growth factor (bFGF) (MilliporeSigma).

### Cell Viability Assays

Cells were seeded on laminin-coated 96-well plates at a density of 2,500–5,000 cells per well and incubated overnight at 37°C to allow attachment. The following day, cells were treated with the indicated concentrations of drugs (QST: 0, 5, or 10 nM; AZD1390: 0, 10, or 30 nM). Quisinostat was dissolved in dimethyl sulfoxide (DMSO) and AZD1390 was dissolved in 100% ethanol. Vehicle control wells received DMSO concentration equivalent to that in the wells with the highest concentration of drug. Cells were incubated for 5 days following treatment. For experiments combining IR, GSCs were treated with either 2 Gy or 4 Gy using an X-Rad225XL irradiator (Precision X-Ray, Madison, CT, USA) 2 hours after drug treatment. Cell viability was measured using the CellTiter-Glo (CTG) cell viability assay according to the manufacturer’s instructions (Promega). Luminescence was recorded using a Tecan Spark plate reader. Luminescence values were normalized to vehicle-treated controls. Results represent the mean of at least three independent biological replicates.

### Immunoblot Assays

Cells were treated as indicated and lysed 24 hours later in radioimmunoprecipitation assay (RIPA) buffer supplemented with phosphatase and protease inhibitor cocktails (Thermo Fisher Scientific). Homogenized samples were subsequently incubated for 20 minutes and centrifuged at 15000 rpm for 10 minutes at 4°C. Samples were stored at −80^°^ C until further analysis. For histone extractions, 0.1 U/ml benzonase (Millipore Sigma) was added to RIPA buffer during lysis. Protein concentrations were determined using the Bradford protein assay (Thermo Fisher Scientific). Equal amounts of protein were separated by SDS–PAGE using 3–8% gradient gels for high–molecular weight proteins and 4–12% gradient gels for other targets, followed by transfer to PVDF membranes (MilliporeSigma). Membranes were blocked in 5% nonfat dry milk prepared in Tris-buffered saline containing 0.1% Tween-20 (TBST) for 1 hour at room temperature and incubated with primary antibodies overnight at 4 °C, followed by incubation with the appropriate secondary antibodies. Protein bands were detected either by fluorescence using a Li-COR Odyssey CLx imaging system or by chemiluminescence using a Li-COR Odyssey Fc imaging system (Li-COR Biosciences). Band intensities were quantified using FIJI (ImageJ). Antibodies and dilution information are listed in Table 1.

**Table 1.** Antibodies used in immunoblot experiments.

| **Antibody** | **Vendor** | **Catalog #** | **Dilution** | **RRID** |
|---|---|---|---|---|
| Phospho-ATM (S1981) | Abcam | ab81292 | 1:2000 | AB_1640207 |
| ATM | Abcam | ab78 | 1:500 | AB_306089 |
| γH2AX (S139) | Abcam | ab11174 | 1:2000 | AB_297813 |
| Phospho-KAP1 (Ser824) | Bethyl | A300-767A | 1:1000 | AB_669740 |
| Phospho-KAP1 (S824) | Abcam | ab70369 | 1:1000 | AB_1209417 |
| Phospho-RAD50 (Ser635) | Cell Signaling Technology | 14223 | 1:1000 | AB_279843 |
| Phospho-RAD50 (Ser635) | Cell Signaling Technology | 74778 | 1:1000 | AB_3740842 |
| Acetylated histone H3 (Lys9/Lys14) | Cell Signaling Technology | 9677 | 1:1000 | AB_114765 |
| Histone H3 | Cell Signaling Technology | 14269S 1 | 1:1000 | AB_2756816 |
| β-actin | Bio-rad | MCA5775GA | 1:5000 | AB_2571580 |
| Vinculin | Sigma, Millipore | V9131 | 1:10000 | AB_477629 |
| Goat anti-Rabbit IgG, DyLight™ 800 | Thermo Fisher Scientific | SA5-10036 | 1:10000 | AB_2556616 |
| Goat anti-Rabbit IgG, DyLight™ 600 | Thermo Fisher Scientific | 35519 | 1:10000 | AB_1965956 |
| F(ab')2-Goat anti-Mouse IgG, HRP | Thermo Fisher Scientific | A24524 | 1:10000 | AB_2535993 |
| F(ab')2-Goat anti-Rabbit IgG, HRP | Thermo Fisher Scientific | A24537 | 1:10000 | AB_2536005 |

### RNA-seq analysis

Tissue collection and detailed method for RNA-seq analysis for *in vivo* tissues are described in previous study (10). For *in vitro* RNA-seq analysis, cells were seeded on P100 tissue culture dishes and were allowed to attach overnight. Following attachment, GB126 cells were incubated with either DMSO, 25nM QST, 4 Gy of IR or 25nM QST and 4 Gy of IR (with IR treatment performed 1hr after QST treatment) for 72 hrs. After 72 hrs, RNA was isolated and quantified for further analysis. The extracted RNA was used for library preparation using the QIAseq FastSelect RNA Lib HMR Kit (Cat# 334235, Qiagen) according to the manufacturer’s instructions. Libraries were quantified using a Qubit fluorometer, and quality and fragment size distribution were evaluated using QIAxcel. Libraries were pooled and sequenced on an Illumina NovaSeq X Plus platform using paired-end reads. After the sequencing, raw reads were processed with fastp for quality filtering and adapter trimming. Trimmed reads were aligned using the STAR aligner to the human reference genome (hg38). Alignment and quality metrics were summarized using MultiQC. Lowly expressed genes were filtered using filterByExpr, and normalization was performed using the TMM method in edgeR (v4.0.16). Differential gene expression analysis was performed using the limma R package (v3.58.11) (15). Linear models were fitted to normalized expression values with *lmFit*, followed by empirical Bayes moderation using *eBayes*. Genes with FDR q < 0.05 were considered significant. Gene set enrichment analysis (GSEA) was performed using GSEA software from MSigDB with gene sets of 10-500 genes, using the classic scoring scheme (16). Pathways with FDR q < 0.05 were considered significant.

### Orthotopic xenograft glioma survival study

All *in vivo* experiments were performed in accordance with the guidelines of St. Joseph’s Hospital and Medical Center and Barrow Neurological Institute under a protocol approved by the Institutional Animal Care and Use Committee (IACUC). Animals were maintained on a 12 h light–dark cycle with ad libitum access to food and water. Six-week-old Hsd:Athymic Nude-Foxn1^nu mice (RRID:MGI:5652489; The Jackson Laboratory) were used for orthotopic implantation of luciferase-expressing GB126 cells. Mice were anesthetized with isoflurane, and 2 µL of cell suspension (50,000 cells/µL) was stereotactically injected into the right striatum 2 mm lateral to the bregma and 2.5 mm ventral. Tumor growth was monitored by bioluminescence imaging, and treatment was initiated once the *In Vivo* Imaging System (IVIS) signal reached the exponential growth phase. Mice were randomized into treatment groups (n = 10 per group). QST (10 mg/kg) and AZD1390 (15 mg/kg) were administered intraperitoneally. QST was administered three times per week (Monday, Wednesday, and Friday) throughout the study. AZD1390 was administered 1 hour after QST, followed by 0.6 Gy IR delivered using an X-Rad225XL irradiator (Precision X-Ray, Madison, CT, USA). AZD1390 and irradiation were administered daily for five consecutive days for one week only. Mice were monitored daily and euthanized upon the onset of neurological symptoms or when moribund.

### Data analysis and statistics

Statistical analyses were performed using GraphPad Prism (version 10). IC_50_ values were calculated using the log(inhibitor) vs. response (variable slope) nonlinear regression model. Drug interaction effects were evaluated using the Bliss independence model implemented in SynergyFinder Plus. Data are presented as mean and ±SD of at least 3 biological replicates. Significance of differences between treatment conditions or cell lines was tested with 2-way ANOVA with Tukey’s multiple comparisons test. Survival studies were analyzed using the Kaplan-Meier method with the Mantel-Cox log-rank test. Statistical significance was defined as *P* less than 0.05.

## Results

### Quisinostat downregulates transcription of homologous recombination pathway components

In our previous work, we demonstrated that HDAC inhibitor QST is cytotoxic to human GSCs and serves as a potent radiosensitizer in an orthotopic PDX model of GBM (10). Although QST combined with IR prolonged survival *in vivo*, tumors ultimately progressed. Acute treatment studies from that work further demonstrated elevated levels of DNA DSBs in tumors treated with QST and IR combination (QST+IR) (10), consistent with therapy-induced DNA damage that was not fully resolved. To extend these findings and more broadly characterize the transcriptional response to QST and IR, we performed further analysis of the RNA-sequencing (RNA-seq) dataset generated from orthotopic tumors in Lo Cascio et al (10). Gene set enrichment analysis (GSEA) of the top 500 gene sets revealed broad transcriptional changes affecting neuronal differentiation, cell cycle regulation, and DNA damage response pathways. Among these, homologous recombination (HR) emerged as one of the most significant suppressed pathways (Fig. 1a). Given the established role of HR as a primary determinant of DSB repair fidelity in GBM (12), we next asked whether QST+IR exerted a comparable transcriptional effect in patient-derived GSCs *in vitro*. To this end, we performed RNA-seq analysis on GSCs treated with QST and IR *in vitro*. Enrichment mapping of the top 450 gene sets revealed convergent pathway alterations, with HR again appearing as a strongly suppressed pathway cluster (Fig. 1b). Waterfall plots confirmed widespread downregulation of HR-associated transcripts both *in vivo* (Fig. 1c) and *in vitro* (Fig. 1d), with multiple genes reaching statistical significance (p < 0.05 to p < 0.0001). Collectively, these findings demonstrate that QST and IR coordinately suppress the HR transcriptional program in both orthotopic tumors and *in vitro* patient-derived GSC cultures.

**Figure 1.**
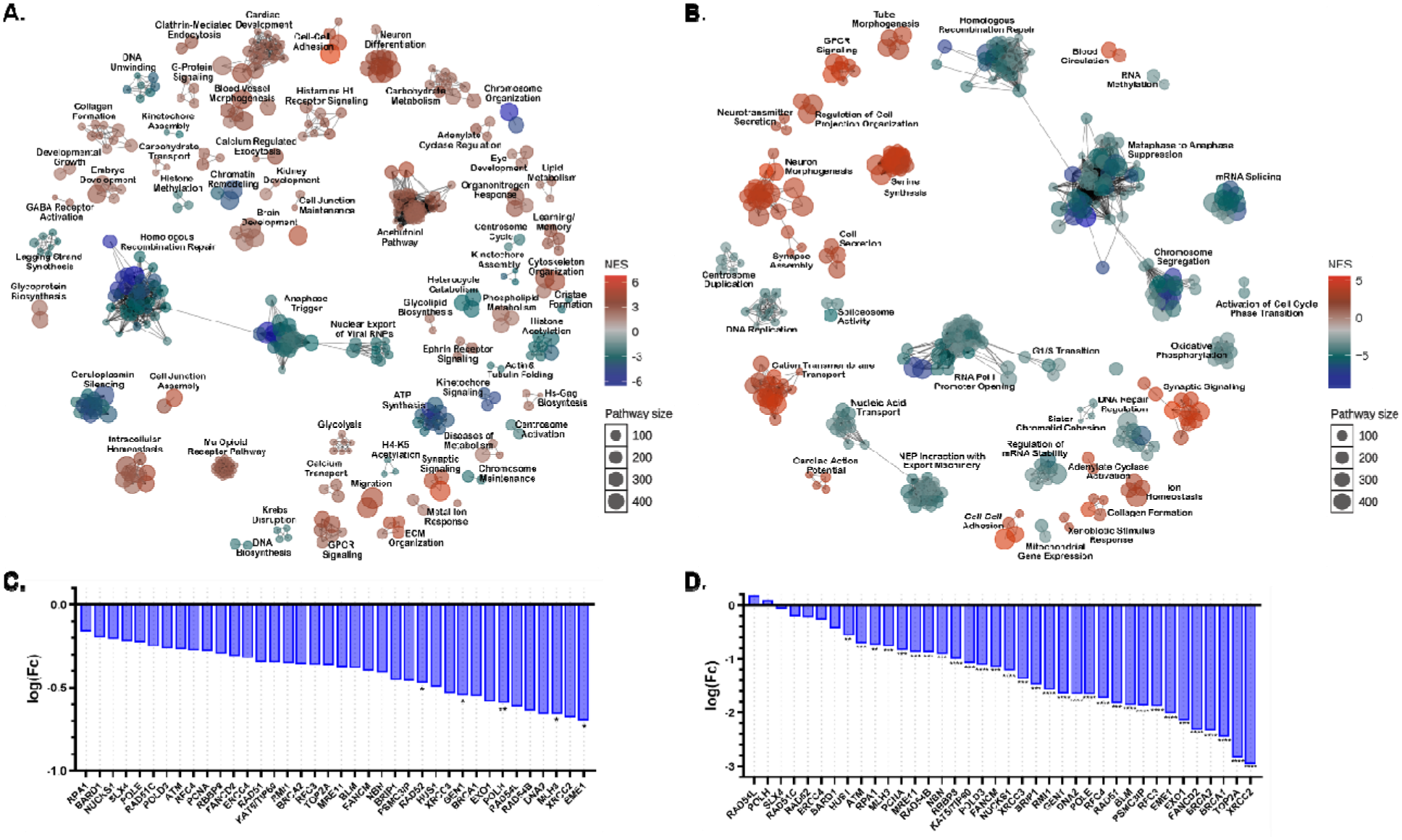
QST combined with radiation suppresses homologous recombination programs *in vitro* and *in vivo*. **(a)** Enrichment map of RNA-seq data from endpoint orthotopic PDX tumors (GB126) comparing vehicle- and QST + IR-treated groups, showing significantly enriched pathways among the top 500 gene sets, colored by normalized enrichment score (NES). **(b)** Enrichment map of RNA-seq data from GB126 GSCs treated with QST + IR *in vitro*, showing significantly enriched pathways among the top 450 gene sets. **(c)** Waterfall plot showing differential expression of homologous recombination pathway genes *in vivo* derived from the dataset in **(a)**. **(d)** Waterfall plot showing differential expression of homologous recombination genes *in vitro* derived from the dataset in **(b)**. \**P*<0.05; \*\**P*<0.01; \*\*\**P*<0.001; \*\*\*\**P*<0.0001.

### Combined inhibition of HDAC and ATM enhances radiation sensitivity in GSCs

ATM is a central kinase governing HR-mediated DSB repair and a key inducer of the radiation-induced DNA damage response (14, 17). Given the broad suppression of HR transcription by QST and IR, we reasoned that pharmacological inhibition of ATM would further impair DSB repair capacity and potentiate the therapeutic efficacy of QST and IR. To test this hypothesis, we evaluated the anti-proliferative effects of combining QST with the brain-penetrant ATM inhibitor, AZD1390 (18, 19), across five patient-derived GSC lines. Cells were treated with escalating concentrations of QST and AZD in the presence or absence of IR, and cell viability was assessed five days later using the CellTiter-Glo luminescent assay (Fig. 2, S1). In GB126 cells, single-agent QST and AZD1390 each reduced cell viability in a dose-dependent manner. The addition of 4 Gy of IR further diminished viability compared with either drug alone, and the triple combination of QST, AZD, and IR produced reductions that exceeded those observed with any dual treatment combinations (Fig. 2a, c). A comparable pattern was observed in the GB239 and GBM43 cell lines, where the triple combination at 2 Gy IR markedly suppressed cell viability relative to IR alone or to either inhibitor paired with IR (Fig. 2b, d and Fig. S1a, d). There were two lines, GB282 and GB187, that did not have a synergistic response (Fig. S1b-f), suggesting that response to the combination of AZD, QST, and IR is not universal and can vary based on other genetic aberrations.

**Figure 2.**
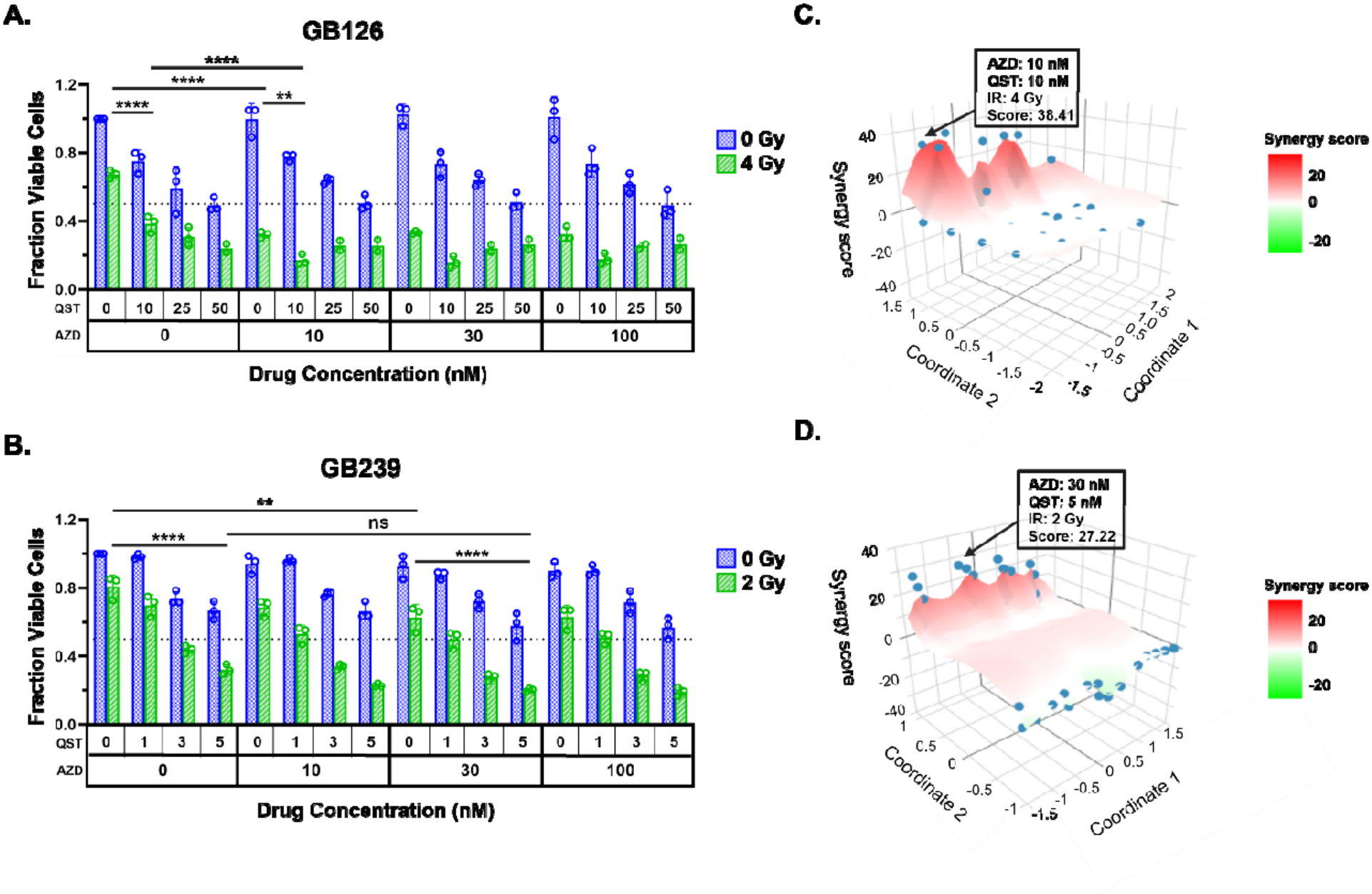
Combined HDAC and ATM inhibition enhances radiation sensitivity in patient-derived GSCs. **(a–b)** Cell viability of GSCs treated with indicated concentrations of QST and AZD1390 in the presence or absence of ionizing radiation (IR; 2 Gy or 4 Gy as indicated). Viability was measured 5 days after treatment by CellTiter-Glo. Data represent mean ± SD from three independent biological replicates. **(c-d)** Bliss synergy landscapes showing interaction between QST and AZD under radiation conditions in the corresponding GSC lines. Bliss scores >10 indicate synergistic interactions. Two-way ANOVA analysis performed on cell viability data at the indicated synergistic dose combinations. \**P*<0.05; \*\**P*<0.01; \*\*\**P*<0.001; \*\*\*\**P*<0.0001.

To quantify the drug interactions among these agents, we applied the Bliss independence model. Synergy landscapes for GB126 revealed a peak Bliss score of 38.41 at AZD1390 10 nM, QST 10 nM, and 4 Gy IR (Fig. 2c). In GB239, a peak Bliss score of 27.22 was observed at AZD 30 nM, QST 5 nM, and 2 Gy IR (Fig. 2d). These scores exceeded the commonly used synergy threshold of >10. Cell viability at these synergistic concentrations was significantly reduced by the triple combination relative to other conditions (*P* < 0.01–0.0001). Collectively, these data demonstrate that simultaneous inhibition of HDAC and ATM produces synergistic radio-sensitization across multiple GBM patient-derived lines.

### Triple combination therapy disrupts ATM signaling, augments histone acetylation, and drives persistent DNA damage accumulation in GSCs

To define the mechanistic basis of the observed synergy, we performed immunoblot analysis on GB126 GSCs treated with synergistic concentrations established in Figure 2c. Whole-cell protein lysates were harvested 24 hours post-treatment and probed for markers of ATM pathway activity, histone acetylation, and DNA damage (Fig. 3a).

**Figure 3.**
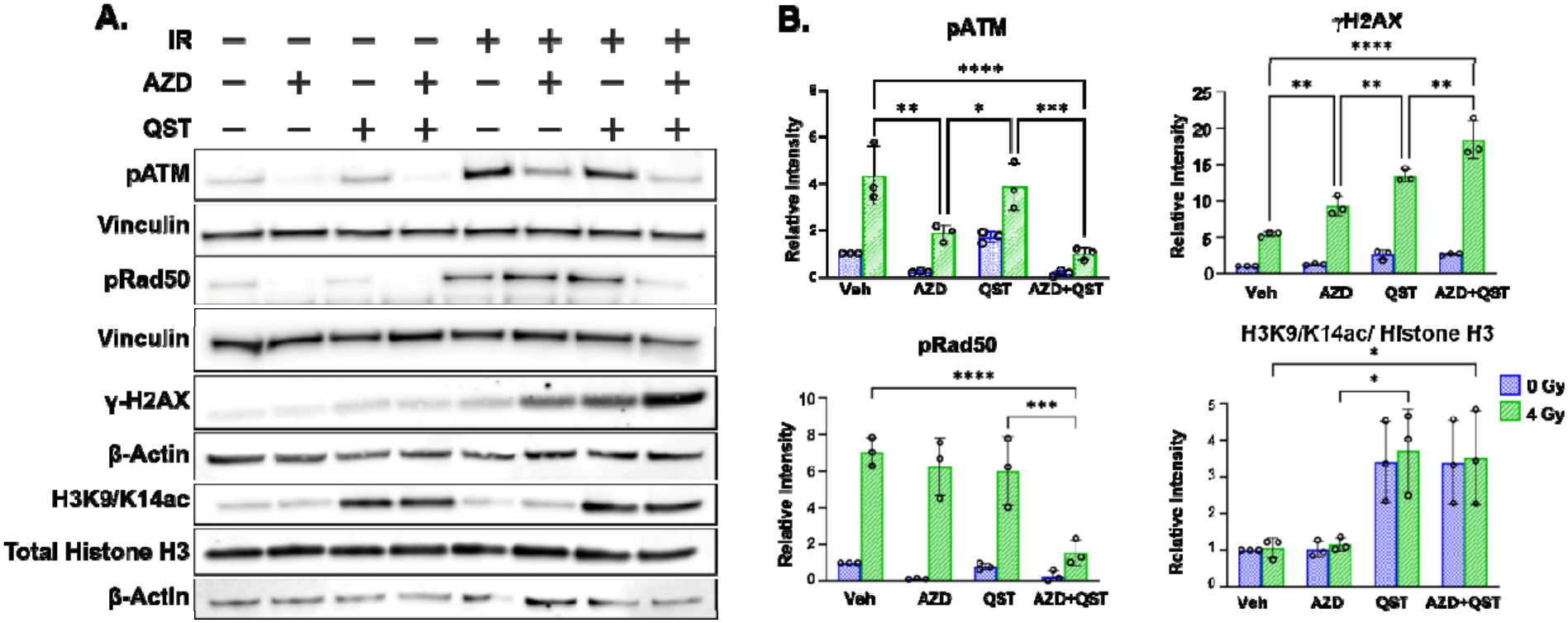
AZD1390, QST, and radiation cooperatively suppress ATM-mediated DDR signaling and increase DNA damage markers in GSCs. **(a)** Immunoblot analysis of proteins targeted by ATM, HDAC, and DNA damage response activation in GB126 GSCs treated with AZD1390 (10 nM), QST (10 nM), and ionizing radiation (IR; 4 Gy) alone or in combination. Cells were harvested 24 hours after treatment and probed for pATM, pRAD50, acetylated histone H3, total histone H3, and γH2AX. **(b)** Quantification of band intensities from **(a)**. Data represent mean ± SD from three independent biological replicates. *P<0.05; **P<0.01; ***P<0.001; ****P<0.0001. Uncropped original blots have been shown in supplementary figure 5A & B.

Consistent with the mechanism of action of AZD1390, treatment with AZD alone, AZD + IR or QST + IR significantly decreased phosphorylation of ATM (S1981), Rad50 (S635), and KAP1 (S824) relative to vehicle controls (Fig. 3, S2) QST alone did not significantly alter ATM pathway signaling, as expected given that HDAC inhibition does not directly target ATM kinase activity. In contrast, QST robustly increased histone H3 acetylation (H3K9/K14ac), confirming on-target HDAC engagement, whereas IR and AZD1390 had no discernible effect on acetylation (Fig. 3). Notably, γH2AX levels, an established marker of unresolved DNA DSBs, were significantly elevated following QST + IR and AZD + IR treatment and were highest in the triple combination group (QST + AZD + IR) (Fig. 3). Similar patterns of protein expression were seen when GB239 was treated with the synergistic combination shown in Figure 2d, although statistically significant changes were not seen with γH2AX (Fig. S3).

This finding is consistent with RNA-seq-identified suppression of HR pathway genes (Fig. 1), suggesting that impaired transcription of HR components contributes to the persistence of DNA damage and the observed synergy. Together, these results indicate that QST and AZD act through complementary and non-overlapping mechanisms to suppress ATM-dependent DNA repair signaling and amplify DNA damage accumulation in GBM cells, providing a mechanistic rationale for their combination with radiotherapy.

### Addition of ATM inhibition to QST and radiotherapy fails to further extend survival in an orthotopic GBM PDX model

To determine whether the *in vitro* synergy observed between QST, AZD1390, and IR translated to improved therapeutic outcomes *in vivo*, we employed an orthotopic GB126 PDX model. Tumor-bearing mice were randomized and treated with QST (10 mg/kg), AZD1390 (15 mg/kg), and fractionated IR (0.6 Gy per fraction × 5) as single agents or in combination, as detailed in Fig. 4a. Animal weights were tracked throughout treatment (Fig. S4) to monitor drug toxicity.

**Figure 4.**
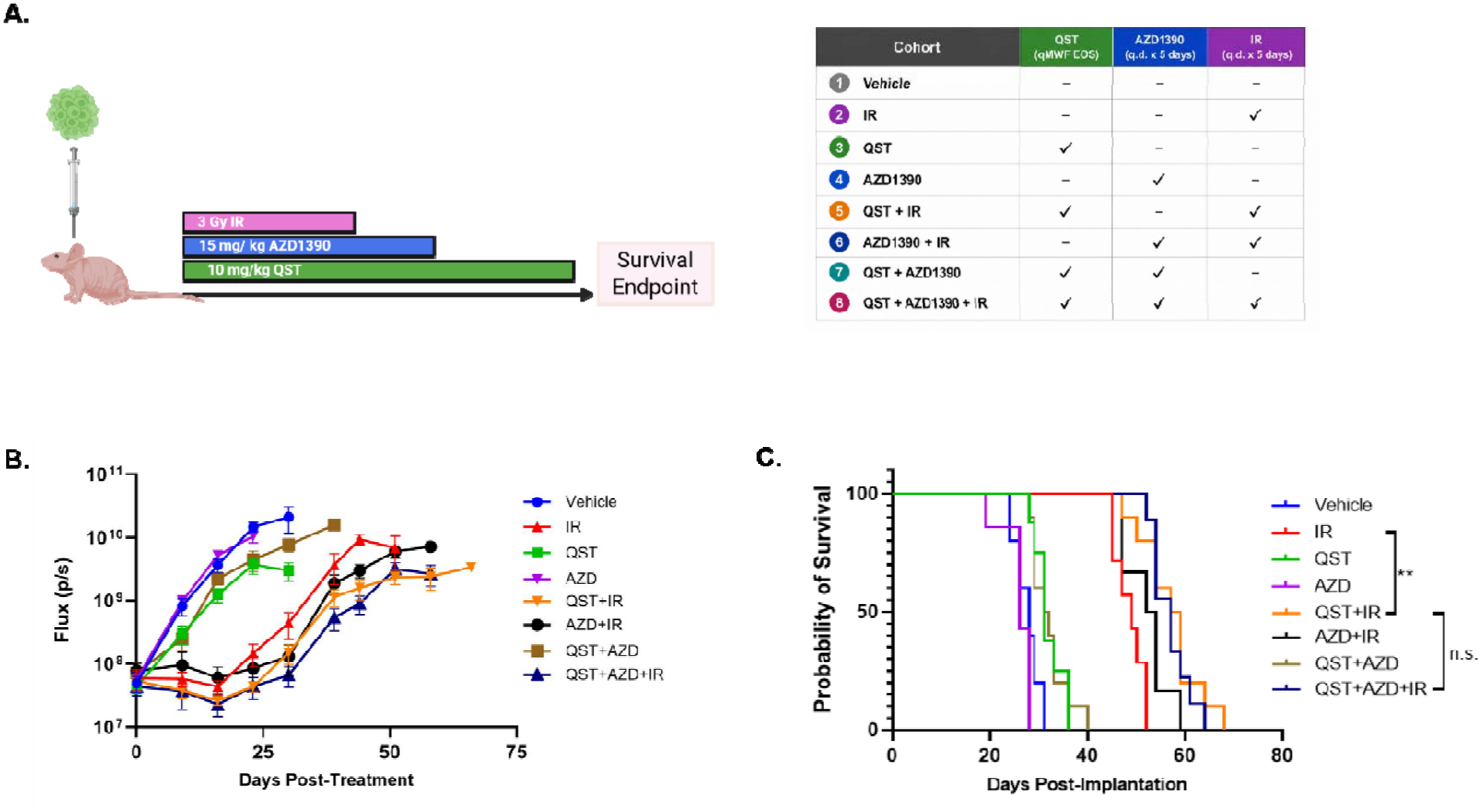
*In vivo* evaluation of QST, AZD1390, and radiation in an orthotopic GBM model. **(a)** Schematic and dosing regimen of the intrarcranial study in GB126 PDX tumors. **(b)** Tumor burden measured by bioluminescence imaging and reported as mean photon flux (p/s) over time for each treatment cohort. **(c)** Kaplan–Meier survival analysis of mice treated with the indicated therapies. *n* = 10 mice per group. Survival differences were evaluated by Mantel–Cox log-rank test. n.s., not significant. \**P* < 0.05; \*\**P* < 0.01; \*\*\**P* < 0.001; \*\*\*\**P*<0.0001.

Bioluminescence imaging demonstrated that IR-containing treatment groups showed delayed tumor growth compared to other groups (Fig. 4b). Among the single modality treatment groups, Kaplan–Meier survival analysis confirmed that IR significantly prolonged survival compared with vehicle-treated mice (median survival: 49 vs 28 days, P<0.0001; Fig. 4c, Table 2). Similarly, QST alone modestly improved survival relative to vehicle (median survival: 31 vs 28 days, P*=*0.0071), whereas AZD1390 alone had no significant effect (median survival: 26 vs 28 days, P*=*0.1598).

**Table 2.** Pairwise log-rank comparisons of survival across treatment groups.

| **Treatment Groups** | **Median Survival Group 1 (Days)** | **Median Survival Group 2 (Days)** | **P Value** |
|---|---|---|---|
| Veh vs IR | 28 | 49 | <0.0001 |
| Veh vs QST | 28 | 31 | 0.0071 |
| Veh vs AZD | 28 | 26 | 0.1598 (n.s.*) |
| Veh vs QST+IR | 28 | 58 | <0.0001 |
| Veh vs AZD+IR | 28 | 53 | 0.0002 |
| Veh vs QST+AZD | 28 | 32 | 0.0040 |
| Veh vs QST+AZD+IR | 28 | 57 | <0.0001 |
| IR vs QST | 49 | 31 | 0.0002 |
| IR vs AZD | 49 | 26 | 0.0003 |
| IR vs QST+IR | 49 | 58 | 0.0010 |
| IR vs AZD+IR | 49 | 53 | 0.0593 (n.s.) |
| IR vs QST+AZD | 49 | 32 | <0.0001 |
| IR vs QST+AZD+IR | 49 | 57 | 0.0001 |
| QST vs AZD | 31 | 26 | 0.0005 |
| QST vs QST+IR | 31 | 58 | <0.0001 |
| QST vs AZD+IR | 31 | 53 | 0.0004 |
| QST vs QST+AZD | 31 | 32 | 0.9072 (n.s.) |
| QST vs QST+AZD+IR | 31 | 57 | <0.0001 |
| AZD vs QSR+IR | 26 | 58 | <0.0001 |
| AZD vs AZD+IR | 26 | 53 | 0.0007 |
| AZD vs QST+AZD | 26 | 32 | 0.0001 |
| AZD vs QST+AZD+IR | 26 | 57 | <0.0001 |
| QST+IR vs AZD+IR | 58 | 53 | 0.0943 (n.s.) |
| QST+IR vs QST+AZD | 58 | 32 | <0.0001 |
| QST+IR vs QST+AZD+IR | 58 | 57 | 0.5929 (n.s.) |
| AZD+IR vs QST+AZD | 53 | 32 | 0.0002 |
| AZD+IR vs QST+AZD+IR | 53 | 57 | 0.0835 (n.s.) |
| QST+AZD vs QST+AZD+IR | 32 | 57 | <0.0001 |

\*n.s.: not significant

Among combination treatments, QST + IR (median survival: 58 vs 49 days, P=0.0010) and the triple combination QST + AZD + IR (median survival: 57 vs 49 days, P=0.0001) significantly prolonged survival compared with IR alone. In contrast, AZD + IR did not significantly improve survival relative to IR alone (median survival: 53 vs 49 days, P=0.0593). No additional benefit was observed with the addition of AZD to QST + IR, as survival was comparable between QST + IR and QST + AZD + IR groups (P=0.5929).

Together, these findings indicate that despite compelling molecular and cellular evidence for synergy *in vitro*, the combination of QST and AZD1390 at the tested doses does not significantly enhance the efficacy of radiotherapy in the orthotopic GB126 GBM PDX model. This discrepancy may reflect pharmacological limitations within the tumor microenvironment or the need for dose optimization.

## Discussion

Both HDAC inhibition and DDR targeting have emerged as promising strategies to overcome therapeutic resistance in GBM (2, 4, 8, 12, 20–22). Because radiotherapy remains a central component of GBM treatment, combinations that impair DNA repair while enhancing radiation-induced damage are particularly attractive (12, 21, 23, 24). In this study, we found that the brain-penetrant HDAC inhibitor QST (10) and the ATM inhibitor AZD1390 produced synergistic radiosensitization in multiple patient-derived GSC models *in vitro.* Mechanistically, QST combined with IR suppressed homologous recombination (HR)-associated transcriptional programs, while AZD1390 inhibited canonical ATM signaling. The triple combination further increased accumulation of γH2AX, consistent with persistent unresolved DNA damage. However, despite this strong mechanistic and cellular rationale, the addition of AZD1390 did not improve survival beyond QST + IR in an orthotopic GBM model under the dosing conditions tested. Together, these findings highlight both the promise and the limitations of combining epigenetic and DDR-targeted therapies in GBM.

Our findings extend prior work showing that HDAC inhibitors can function as radiosensitizers in GBM by altering chromatin accessibility, transcriptional programs, and DNA repair capacity (9, 20, 25–27). In our previous study, QST demonstrated brain penetration, on-target activity, and significantly prolonged survival in orthotopic GBM models when combined with radiation (10). The current study builds on that work by testing whether additional targeting of ATM, a central kinase in the response to DNA double-strand breaks (14, 17), could further exploit a repair vulnerability created by HDAC inhibition. Suppression of HR-associated genes following QST + IR treatment in both orthotopic tumors and GSC cultures suggests that QST primes GBM cells for greater dependence on DDR signaling. In this context, ATM inhibition was a rational strategy to further constrain repair of radiation- and QST-induced DNA damage.

*In vitro,* QST increased histone acetylation and suppressed HR-associated transcription, whereas AZD1390 inhibited ATM pathway signaling through reduced phosphorylation of ATM, RAD50, and KAP1 (Fig. 3, S2). These data support complementary, rather than redundant action of the two agents. The increase in γH2AX observed with the triple combination is therefore most consistent with impaired resolution of DNA double-strand breaks rather than simply greater initial damage. These findings support a model in which HDAC inhibition weakens transcriptional support for high-fidelity repair, while ATM inhibition blocks acute signaling responses required for damage processing after radiation. This dual pressure likely underlies the strong synergy observed in several GSC models.

Notably, the variable *in vitro* response across GSC models reveals the heterogeneity of GBM and suggests that sensitivity to combined HDAC and ATM inhibition is context dependent rather than uniform across tumors. Although synergy was observed in several models, responses differed between lines, indicating tumor-intrinsic variability in dependence on DDR pathways. This heterogeneity is consistent with the broader GBM literature (22, 28) and argues against a biomarker-agnostic approach to DDR-targeted combinations. Factors including baseline DDR dependency, molecular subtype, and stemness-associated programs may all contribute to therapeutic response.

Prior studies have reported greater radiosensitization with ATM inhibition in p53-mutant GBM models, whereas TP53-wildtype tumors may be less dependent on ATM-mediated survival following radiation (18, 19, 30). In addition, HDAC inhibitors have been reported to influence p53 signaling through regulation of p53 acetylation and downstream transcriptional programs (29). However, the strongest *in vitro* synergy in our study was also observed in TP53-wildtype models, including GB126 and GB239, suggesting that p53 status alone is insufficient to explain response to the triple combination. The heterogeneous nature of GBM introduces additional complexity when translating *in vitro* findings to orthotopic models, where the tumor microenvironment can influence tumor growth, drug distribution, and therapeutic response in ways not captured *in vitro* (2, 12, 22, 31).

The absence of added survival benefit with AZD1390 *in vivo*, despite robust *in vitro* synergy, should be interpreted within the context of the treatment conditions evaluated. As with many oncology combination studies, the balance between tolerability, exposure, and pathway modulation may have limited the ability to realize the cooperative effects seen *in vitro*. The regimen used here was reduced relative to prior AZD1390-radiation studies due to tolerability concerns with the triple combination, potentially limiting effective pathway inhibition *in vivo* (10, 18, 19). More broadly, this reflects a common challenge in oncology combination studies, where tolerability constraints can compromise delivery of pharmacologically effective doses (32–34). In addition, patient-derived GBM models harbor diverse mutational profiles that can influence DDR pathway dependency and treatment susceptibility, potentially contributing to variability in therapeutic outcomes across models (28, 35, 36).

In summary, combined HDAC and ATM inhibition produced strong radiosensitization and mechanistic evidence of impaired DNA damage resolution in patient-derived GSCs *in vitro*; however, this effect did not translate into improved survival beyond QST + IR in an orthotopic p53-wildtype GBM model under the dosing conditions tested. These findings refine the translational landscape for DDR-targeted therapy in GBM by showing that biologically rational combinations require not only mechanistic synergy, but also appropriate pharmacologic exposure for target engagement and genetic context for a response *in vivo*. Further studies will be required to determine the conditions under which HDAC and DDR-targeted inhibitor combinations can achieve durable therapeutic benefit in GBM.

### Conclusions

This study demonstrates that combined inhibition of HDAC and ATM using quisinostat and AZD1390 enhances radiosensitivity in patient-derived GSCs, with synergistic effects observed in three of five GSC models *in vitro*. Despite robust mechanistic and cellular evidence of synergy, ATM inhibitor did not confer additional survival benefit beyond quisinostat plus radiation in an orthotopic GBM model *in vivo* under the dosing conditions evaluated. These findings establish an epigenetic basis for HR vulnerability in GBM and highlight that pharmacological exposure, dosing-schedules and genetic context are critical determinants of *in vivo* efficacy for DDR-targeted combination strategies.

## Data Availability

All data are available within the article and can be accessed from public repositories or in the Supporting Data Values file.

## Ethics approval

Human surgical GBM specimens used for deriving patient-derived glioma stem cells and xenograft models were collected from the Ivy Brain Tumor Center at St. Joseph’s Hospital and Barrow Neurological Institute under protocol approved by WCG Institutional Review Board (No. 20200850). The samples were deidentified and conformed to the Institutional Review Board’s protocol. Animal husbandry was performed in accordance with the guidelines of St. Joseph’s Hospital and Medical Center and Barrow Neurological Institute under the protocol approved by the Institutional Animal Care and Use Committee.

## Author contributions

SM, AT, ACT, NS, GU, SP, and SQH conceived and designed experiments. SQH, KC, EC, TD, LS, KO, and NES performed *in vitro* experiments, and analyzed the data. GU, SP, AT, and SM designed *in vivo* experiments and GU coordinated and oversaw all *in vivo* experiments. GU, EH, and KKW performed all *in vivo* experiments. SD performed RNA-seq library preparation for the *in vitro* RNA-seq samples. ACT and SD contributed to interpretation of RNA-seq results. GU, SP, and SM edited the manuscript. All authors reviewed the manuscript. SM coordinated and supervised the project. SQH and SM wrote the manuscript.

## Supporting information

Manuscript

## Acknowledgements

We thank the entire Mehta lab for helpful discussions and technical assistance including Zackariah Brownell for assistance with *in vivo* experiments. We thank VUGENE for assistance with bioinformatics, analysis of RNA-seq data, and generation of the enrichment maps in Figure 1. Schematic in Figure 4 was generated using BioRender.com.

## Abbreviations

GBM: Glioblastoma
BBB: Blood-brain barrier
HDAC: Histone deacetylase
DDR: DNA damage response
HDACi: HDAC inhibitor
QST: Quisinostat
GSC: Glioma stem cell
ATM: Ataxia-telangiectasia mutated
DSB: Double-strand break
IR: Ionizing radiation
PDX: Patient-derived xenograft
EGF: Epidermal growth factor
bFGF: Basic fibroblast growth factor
DMSO: Dimethyl sulfoxide
CTG: CellTiter-Glo
RIPA: Radioimmunoprecipitation assay
TBST: Tris-buffered saline containing tween-20
GSEA: Gene set enrichment analysis
IACUC: Institutional animal care and use committee
RNA-seq: RNA sequencing
HR: Homologous recombination
TMZ: Temozolomide
MGMT: O6-methylguanine-DNA methyltransferase

## References

1. Price M, Ballard CAP, Benedetti JR, Kruchko C, Barnholtz-Sloan JS, Ostrom QT. CBTRUS Statistical Report: Primary Brain and Other Central Nervous System Tumors Diagnosed in the United States in 2018-2022. Neuro Oncol. 2025;27(Supplement_4):iv1–iv66.

2. Singh S, Dey D, Barik D, Mohapatra I, Kim S, Sharma M, et al. Glioblastoma at the crossroads: current understanding and future therapeutic horizons. Signal Transduct Target Ther. 2025;10(1):213.

3. Stupp R, Mason WP, van den Bent MJ, Weller M, Fisher B, Taphoorn MJ, et al. Radiotherapy plus concomitant and adjuvant temozolomide for glioblastoma. N Engl J Med. 2005;352(10):987–96.

4. Tehrani GA, Kubick RN, Zarodniuk M, Datta M. Histone deacetylase inhibitors sensitize glioblastoma models to temozolomide and reprogram immunosuppressive myeloid cells. Scientific Reports. 2025;15(1):36804.

5. Cheng Y, He C, Wang M, Ma X, Mo F, Yang S, et al. Targeting epigenetic regulators for cancer therapy: mechanisms and advances in clinical trials. Signal Transduct Target Ther. 2019;4:62.

6. Li G, Tian Y, Zhu W-G. The Roles of Histone Deacetylases and Their Inhibitors in Cancer Therapy. Frontiers in Cell and Developmental Biology. 2020;Volume 8 - 2020.

7. Lee JH, Choy ML, Ngo L, Foster SS, Marks PA. Histone deacetylase inhibitor induces DNA damage, which normal but not transformed cells can repair. Proc Natl Acad Sci U S A. 2010;107(33):14639–44.

8. Eckschlager T, Plch J, Stiborova M, Hrabeta J. Histone Deacetylase Inhibitors as Anticancer Drugs. Int J Mol Sci. 2017;18(7).

9. Lo Cascio C, McNamara JB, Melendez EL, Lewis EM, Dufault ME, Sanai N, et al. Nonredundant, isoform-specific roles of HDAC1 in glioma stem cells. JCI Insight. 2021;6(17).

10. Lo Cascio C, Margaryan T, Luna-Melendez E, McNamara JB, White CI, Knight W, et al. Quisinostat is a brain-penetrant radiosensitizer in glioblastoma. JCI Insight. 2023;8(22).

11. Singh N, Miner A, Hennis L, Mittal S. Mechanisms of temozolomide resistance in glioblastoma - a comprehensive review. Cancer Drug Resist. 2021;4(1):17–43.

12. Majd NK, Yap TA, Koul D, Balasubramaniyan V, Li X, Khan S, et al. The promise of DNA damage response inhibitors for the treatment of glioblastoma. Neurooncol Adv. 2021;3(1):vdab015.

13. Jackson SP, Bartek J. The DNA-damage response in human biology and disease. Nature. 2009;461(7267):1071–8.

14. Blackford AN, Jackson SP. ATM, ATR, and DNA-PK: the trinity at the heart of the DNA damage response. Molecular cell. 2017;66(6):801–17.

15. Ritchie ME, Phipson B, Wu D, Hu Y, Law CW, Shi W, Smyth GK. limma powers differential expression analyses for RNA-sequencing and microarray studies. Nucleic Acids Res. 2015;43(7):e47.

16. Wu T, Hu E, Xu S, Chen M, Guo P, Dai Z, et al. clusterProfiler 4.0: A universal enrichment tool for interpreting omics data. Innovation (Camb). 2021;2(3):100141.

17. Golding SE, Rosenberg E, Adams BR, Wignarajah S, Beckta JM, O’Connor MJ, Valerie K. Dynamic inhibition of ATM kinase provides a strategy for glioblastoma multiforme radiosensitization and growth control. Cell Cycle. 2012;11(6):1167–73.

18. Durant ST, Zheng L, Wang Y, Chen K, Zhang L, Zhang T, et al. The brain-penetrant clinical ATM inhibitor AZD1390 radiosensitizes and improves survival of preclinical brain tumor models. Sci Adv. 2018;4(6):eaat1719.

19. Chen J, Laverty DJ, Talele S, Bale A, Carlson BL, Porath KA, et al. Aberrant ATM signaling and homology-directed DNA repair as a vulnerability of p53-mutant GBM to AZD1390-mediated radiosensitization. Sci Transl Med. 2024;16(734):eadj5962.

20. Pai P, Das I, Reddy Y, Venkidesh BS, Bhandari P, Madalageri M, et al. Targeting glioblastoma with HDAC inhibitors: insights into hydroxamic acid-based therapeutic strategies. Acta Neuropathologica Communications. 2025;14(1):9.

21. Ferri A, Stagni V, Barila D. Targeting the DNA Damage Response to Overcome Cancer Drug Resistance in Glioblastoma. Int J Mol Sci. 2020;21(14).

22. Dewdney B, Jenkins MR, Best SA, Freytag S, Prasad K, Holst J, et al. From signalling pathways to targeted therapies: unravelling glioblastoma’s secrets and harnessing two decades of progress. Signal Transduct Target Ther. 2023;8(1):400.

23. Drew Y, Zenke FT, Curtin NJ. DNA damage response inhibitors in cancer therapy: lessons from the past, current status and future implications. Nature Reviews Drug Discovery. 2025;24(1):19–39.

24. Zhang J, FG Stevens M, D Bradshaw T. Temozolomide: mechanisms of action, repair and resistance. Current molecular pharmacology. 2012;5(1):102–14.

25. Zhang Y, Wang H, Zhan Z, Gan L, Bai O. Mechanisms of HDACs in cancer development. Front Immunol. 2025;16:1529239.

26. Bezecny P. Histone deacetylase inhibitors in glioblastoma: pre-clinical and clinical experience. Medical Oncology. 2014;31(6):985.

27. Chen R, Zhang M, Zhou Y, Guo W, Yi M, Zhang Z, et al. The application of histone deacetylases inhibitors in glioblastoma. J Exp Clin Cancer Res. 2020;39(1):138.

28. Ellis HP, Greenslade M, Powell B, Spiteri I, Sottoriva A, Kurian KM. Current Challenges in Glioblastoma: Intratumour Heterogeneity, Residual Disease, and Models to Predict Disease Recurrence. Front Oncol. 2015;5:251.

29. Diss E, Nalabothula N, Nguyen D, Chang E, Kwok Y, Carrier F. Vorinostat(SAHA) Promotes Hyper-Radiosensitivity in Wild Type p53 Human Glioblastoma Cells. J Clin Oncol Res. 2014;2(1).

30. Biddlestone-Thorpe L, Sajjad M, Rosenberg E, Beckta JM, Valerie NC, Tokarz M, et al. ATM kinase inhibition preferentially sensitizes p53-mutant glioma to ionizing radiation. Clin Cancer Res. 2013;19(12):3189–200.

31. Silverbush D, Jurgensen L, Freeburg NF, Pooley CS, Boniolo F, Gaiti F, et al. A cellular epigenetic classification system for glioblastoma. Neuro Oncol. 2026.

32. Benada J, Bulanova D, Azzoni V, Petrosius V, Ghazanfar S, Wennerberg K, Sorensen CS. Synthetic lethal interaction between WEE1 and PKMYT1 is a target for multiple low-dose treatment of high-grade serous ovarian carcinoma. NAR Cancer. 2023;5(3):zcad029.

33. Kong A, Mehanna H. WEE1 Inhibitor: Clinical Development. Curr Oncol Rep. 2021;23(9):107.

34. Martorana F, Da Silva LA, Sessa C, Colombo I. Everything Comes with a Price: The Toxicity Profile of DNA-Damage Response Targeting Agents. Cancers (Basel). 2022;14(4).

35. Fomchenko EI, Holland EC. Mouse models of brain tumors and their applications in preclinical trials. Clin Cancer Res. 2006;12(18):5288–97.

36. Urzi O, Gasparro R, Costanzo E, De Luca A, Giavaresi G, Fontana S, Alessandro R. Three-Dimensional Cell Cultures: The Bridge between In Vitro and In Vivo Models. Int J Mol Sci. 2023;24(15).

