## Supplementary material for "Combined HDAC and ATM inhibition with radiation in preclinical glioblastoma models": Manuscript

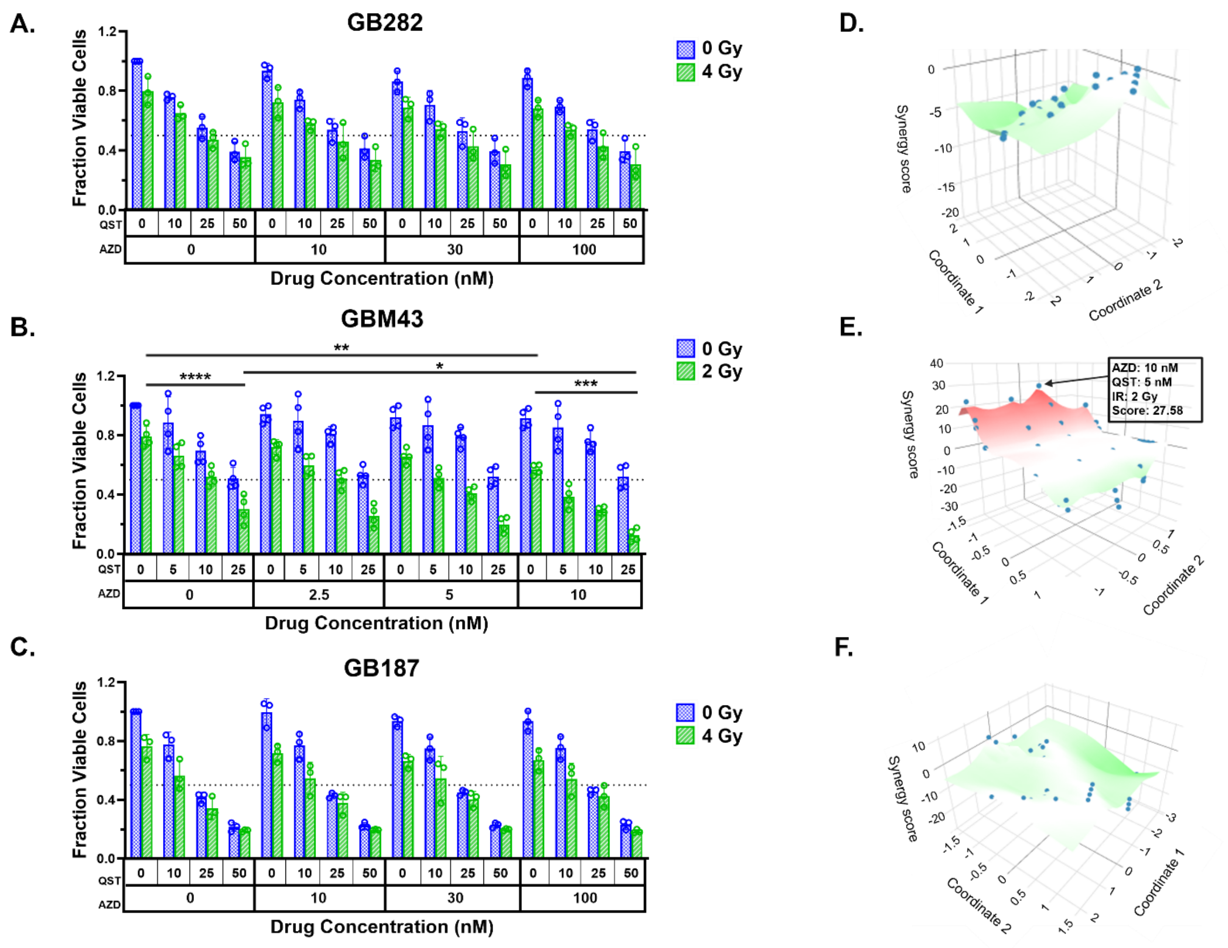


**Supplemental Figure 1. Combined HDAC and ATM inhibition enhances radiation sensitivity in patient-derived GSCs. (A-C)** Cell viability of GSCs treated with indicated concentrations of QST and AZD1390 with or without ionizing radiation. Viability was measured 5 days after treatment by CellTiter-Glo. Data represent mean ± SD from three independent biological replicates. **(D-F)** Bliss synergy landscapes showing interaction between QST and AZD under radiation conditions in the corresponding GSC lines. Bliss scores >10 indicate synergistic interactions, <-10 indicates antagonistic and a score between -10 to 10 indicates additive effects. Two-way ANOVA analysis performed on cell viability data at the indicated synergistic dose combinations. *$P$<0.05; **$P$<0.01; ***$P$<0.001; ****$P$<0.0001.

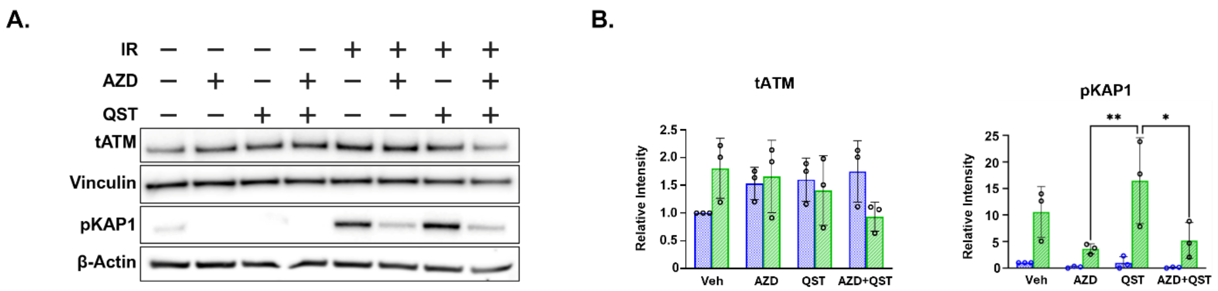


**Supplemental Figure 2. Pharmacodynamic response of AZD1390, QST, and radiation on GSCs *in vitro*. (A)** Immunoblot analysis of homologous recombination-assocociated proteins total ATM and pKAP1 in GB126 GSCs treated with AZD1390 (10 nM), QST (10 nM), and ionizing radiation (IR; 4 Gy) alone or in combination. Cell harvesting and immunoblot methods details discussed in Figure 3. **(B)** Quantification of band intensities from **(A)**. Data represent mean ± SD from three independent biological replicates. *P<0.05. Uncropped original blots have been shown in supplementary figure 5C.

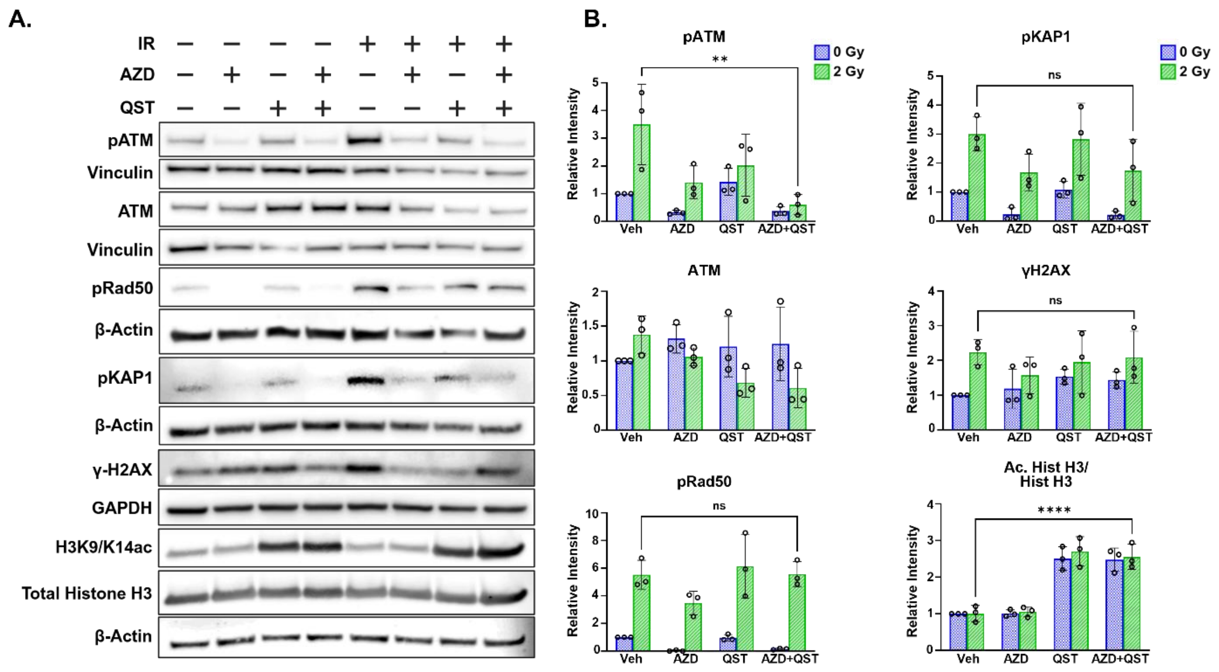


**Supplemental Figure 3. AZD1390, QST, and radiation cooperatively suppress homologous recombination signaling and increase DNA damage in GSCs.**

**(A)** Immunoblot blot analysis of homologous recombination or HDAC inhibitor and DNA damage markers in GB239 GSCs treated with AZD1390 (30 nM), QST (5 nM), and ionizing radiation (IR; 2 Gy) alone or in combination. Cells were harvested 24 hours after treatment and probed for pATM, ATM, pRAD50, pKAP1, γH2AX, acetylated histone H3, and total histone H3. Vinculin, GAPDH, or β-actin were used as loading controls. **(B)** Quantification of band intensities from **(A).** Data represent mean ±SD from four independent biological experiments. * $p < 0.05$, ** $p < 0.01$, ***$p<0.001$, ****$p<0.0001$. Uncropped original blots have been shown in supplementary figure 5D & E.

**A.**

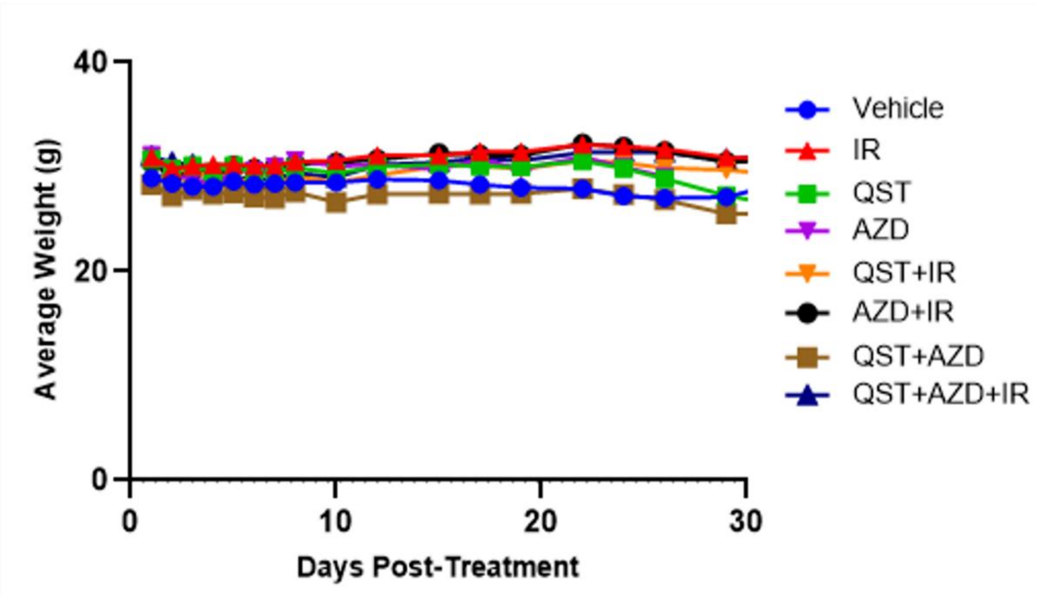


**Supplemental Figure 4. Weights of mice treated with AZD, QST, and IR either alone or in combination are not significantly affected. (A)** Graph depicting the average weight of mice in each treatment cohort over the course of treatment with AZD, QST, and IR.

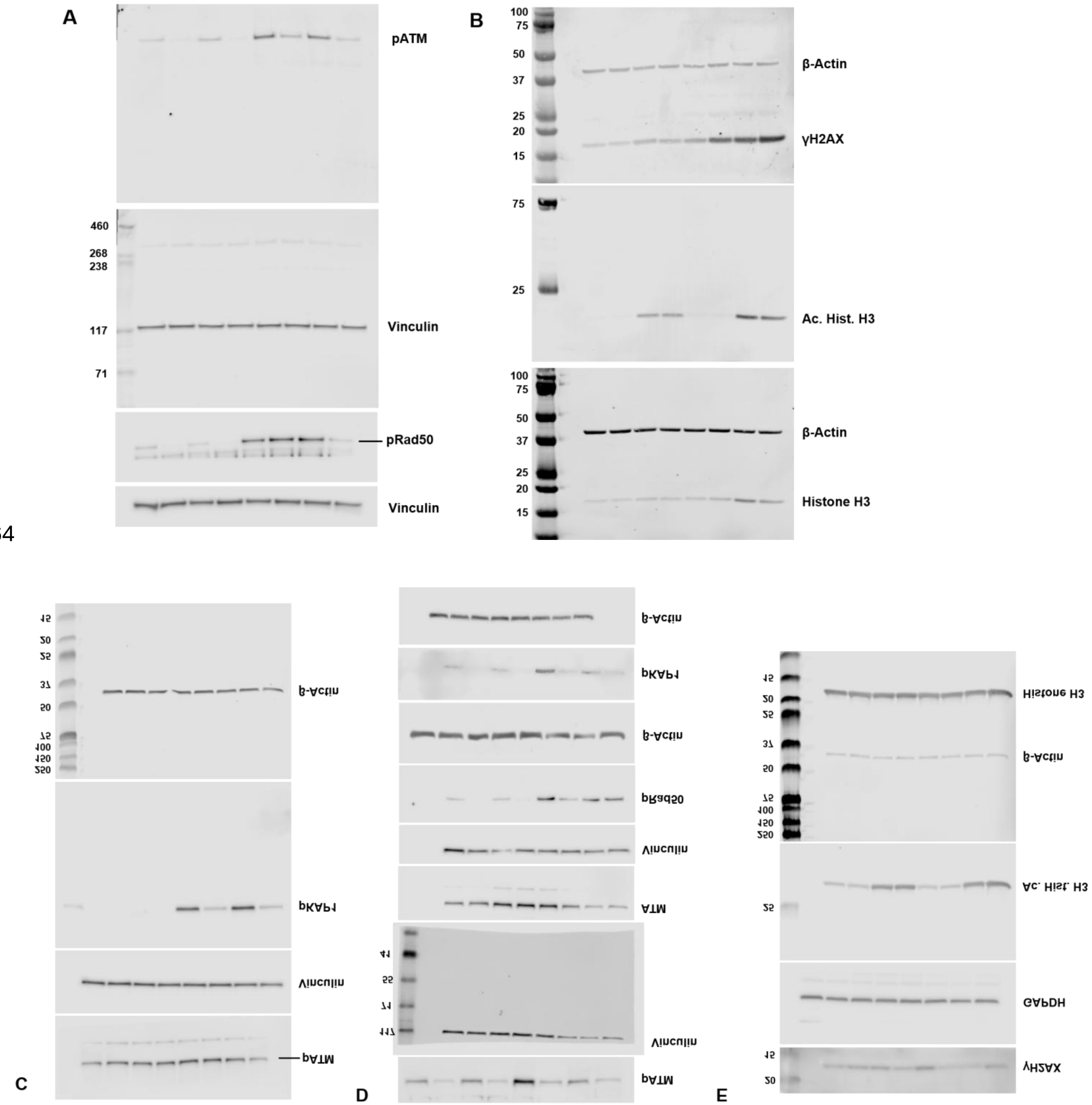


**Supplemental Figure 5. Uncropped original blots:** (A) & (B) Original blots corresponding to Figure 3A, (C) Original blots corresponding to Supplementary Figure 2A, (D) & (E) ) Original blots corresponding to Supplementary Figure 3A
